# Single Cell RNA-Sequence Analyses Reveal Uniquely Expressed Genes and Heterogeneous Immune Cell Involvement in the Rat Model of Intervertebral Disc Degeneration

**DOI:** 10.1101/2022.07.05.498865

**Authors:** Milad Rohanifar, Sade W. Clayton, Garrett Easson, Deepanjali S. Patil, Frank Lee, Liufang Jing, Marcos N. Barcellona, Julie E. Speer, Jordan J. Stivers, Simon Y. Tang, Lori A. Setton

## Abstract

Intervertebral disc (IVD) degeneration is characterized by a loss of cellularity, and changes in cell-mediated activity that drives anatomic changes to IVD structure. In this study, we use single cell RNA-sequencing analysis of cells extracted from the degenerating tissues of the rat IVD following lumbar disc puncture. Two control, uninjured IVDs (L2-3, L3-4) and two degenerated, injured IVDs (L4-5, L5-6) from each animal were examined either at two- and eight-week post-operative time points. The cells from these IVDs were extracted and transcriptionally profiled at a single-cell resolution. Unsupervised cluster analysis revealed the presence of 4 known cell types in both non-degenerative and degenerated IVDs based on previously established gene markers: IVD cells, endothelial cells, myeloid cells, and lymphoid cells. As a majority of cells were associated with the IVD cell cluster, sub-clustering was used to further identify the cell populations of the nucleus pulposus, inner and outer annulus fibrosus. The most notable difference between control and degenerated IVDs was the increase of myeloid and lymphoid cells in degenerated samples at 2- and 8- weeks post-surgery. Differential gene expression analysis revealed multiple distinct cell types from the myeloid and lymphoid lineages, most notably macrophages and B lymphocytes and demonstrated a high degree of immune specificity during degeneration. In addition to the heterogenous infiltrating immune cell populations in the degenerating IVD, the increased number of cells in the AF sub-cluster expressing *Ngf* and *Ngfr*, encoding for p75NTR, suggest that NGF signaling may be one of the key mediators of the IVD crosstalk between immune and neuronal cell populations. These findings provide the basis for future work to understand the involvement of select subsets of non-resident cells in IVD degeneration.

## INTRODUCTION

The intervertebral disc (IVD) provides for motion and flexibility in the spine [1]. Degeneration of the IVD is characterized by a loss of cellularity, changes in composition and loss of hydration that is manifested as changes in disc height and MRI signal intensity. These features of IVD degeneration in the lumbar and cervical spine can contribute to mal-alignment of the adjoining vertebral bodies and affect the integrity and health of the IVD and adjacent neural structures, leaving the IVD vulnerable to progressive damage during the loading conditions of daily living [2–4]. The cell-mediated changes affecting IVD growth, extracellular matrix synthesis and metabolism can affect IVD proteolytic degradation or regeneration, and are of great interest for identifying the relevant cellular and molecular targets to support IVD repair and regeneration.

Cell density of the nucleus pulposus (NP) region are at the lowest in the IVD, and are the principal contributors to loss of cellularity with age. NP cells have long been considered to share features with chondrocytes in the adult IVD, due to their roundedness and high expression of multiple chondrocyte markers including SOX9, type II collagen and aggrecan upon bulk mRNA transcriptional profiling [5–9]. NP cells are derived from the embryonic notochord and express transcriptionally unique markers that reflect their notochordal origins, including CD24, cytokeratins, and the brachyury and FOX transcription factors [10–13]. Cells of the annulus fibrosus (AF) are derived from mesenchyme and express some overlapping and some differing molecular markers from adjacent NP cells. Indeed, there have been numerous studies designed to identify the unique cellular phenotypes of NP and AF cells, as well as non-resident cells involved in degeneration and repair, using tools of bulk RNA transcription or RNA-sequencing, proteomic profiling, flow cytometry and protein assays [5,14–18]. The improved understanding of cell-specific molecular changes is crucial for identifying important protein targets that drive progress degeneration and molecular targets for stem cell mediated IVD repair.

In recent years, single-cell RNA sequencing (scRNA-seq) of IVD cells, both from anatomically distinct regions and for comparisons with pathology, has been used to characterize the phenotype and abundance of distinct cell populations in the IVD in human and animal tissue sources [5,15,17,19–21]. When cells are isolated rapidly and immediately prepared for RNA-sequencing, this technique has potential to reveal the endogenous RNA expression of resident cells and thus identify differences in cell sub-populations in the native IVD [5,15,17,19–22]. Results of sc-RNA-seq from rat and bovine IVD tissues, and from non-degenerate human IVD tissues, rely on mapping cells with similar transcriptomic profiles into “clusters” in order to name and number discrete cell populations in the non-degenerate IVD. While the numbers of clusters is highly variable in the absence of a consensus-based approach, it is most commonly observed that the majority of IVD cells maps to a few clusters based on their expression of common genes (50-99% of isolated cells). Studies have used this approach to identify small but meaningful stem cell or progenitor cell populations in the IVD [15, 20], or to identify relationships between identified clusters and primary cell functions of matrix regulation, stress responses, cell cycle and more [22].

Only a few studies have used sc-RNA-seq to better understand the progression of IVD degeneration at the cellular level. In donor human tissues, sc-RNA-seq has been used to identify genes associated with cells of the degenerative IVD with some findings that corroborate prior work including unique associative expression for CTGF, S100A1/A2, and TNF receptors in cells from degenerated IVDs [5, 22]; both human studies and a study of a rodent bipedal IVD degeneration model also present surprising findings for novel gene involvement that have not yet been confirmed in other studies or species. In some cases, sc-RNA-seq identifies cells that express macrophage markers at high levels [5] consistent with prior literature showing increased involvement of macrophages in animal models and human tissues with IVD degeneration [23]. The IVD is an aneural, alymphatic and avascular structure upon termination of development and growth, however, it can remain immunologically separate from the host over the lifetime of an individual. The presence of macrophage markers in degenerated IVD tissues and cells suggests integrity of the IVD may be disrupted such that cells of the systemic circulation may infiltrate and reside in the IVD. These cells can be expected to be minor and difficult to localize via immunostaining or RT-PCR; for this reason, sc-RNA-seq provides potential to identify the unique phenotypes of these relatively small cell populations in the IVD. For these reasons, we performed scRNA-seq analysis on cells obtained from rat lumbar IVD tissues at two timepoints following induction of IVD degeneration via stab injury [24–28]. In brief, we identified four major cell types in the non-degenerate and degenerative IVD tissues of 2 weeks following surgery, based on the expression of “classical” cell-specific markers for NP and AF cells as identified previously [5]. The cell clustering scheme was held fixed between control and IVD degeneration conditions to estimate the differences in cell numbers and major gene expression levels for respective cell populations within each cluster. Further, we used sub-clustering to identify discrete native cell types within the IVD, and gene set analysis to identify the numerically minor cells of the degenerated IVD and their changes from 2 to 8 weeks after induced IVD degeneration. Our results confirm a set of differentially expressed genes in the degenerated IVD at both timepoints following lumbar disc puncture that point to a role for immune cells, principally macrophages and B-cell subsets that may be infiltrating into the injured IVDs. These findings provide the basis for future work to understand the immune cell specificity in mediating disease progression.

## Materials and Methods

### IVD tissue preparation and single-cell isolation

Rats (n=8, male Sprague-Dawley, 8 weeks-old) underwent surgery for retroperitoneal exposure of the L4-L6 lumbar spine in protocols approved by the Washington University IACUC. The L4-5 and L5-6 IVDs each received a single unilateral stab injury via a 27G needle. The rats were allowed to recover postoperatively for either 2 (n=4) or 8 weeks (n=4) (Figure 1a). Following the respective recovery periods, the animals were euthanized and the IVD tissues were isolated and pooled from L4-5 and L5-6 (LDP= Lumbar disc puncture), or from L2-3 and L3-4 (CON= control), with care to remove peripheral muscle, ligaments, and attachments by microdissection. The isolated IVD tissue was digested in medium containing 0.2% type 2 collagenase (Worthington Biochemical, Lakewood, NK) and 0.3% pronase (Roche) for < 4 h total at 37 °C and 5% CO_2_. After digestion, a cell pellet was obtained by centrifuging the medium for 10 min at 400 rcf. The media was aspirated, and cells were resuspended in PBS (Sigma Aldrich), followed by filtering through a 70 μm filter to get rid of undesired cell debris. The flowthrough was again centrifuged for 10 min at 400 rcf, and the resulting cell pellet was resuspended in PBS. The cells isolated from pooled IVDs for each rat were kept together and considered as a separate sample for a total of n=16 samples in total (n=4 for CON and LDP at 2 weeks post-surgery; n=4 for CON and LDP at 8 weeks post-surgery). Samples of IVD cells were immediately transported to the sequencing facility (Genome Technology and Access Center, Washington University). More than 80% of cells were determined to be viable. All analyses described below were performed on individual rat cell samples unless otherwise indicated.

**Figure 1.**
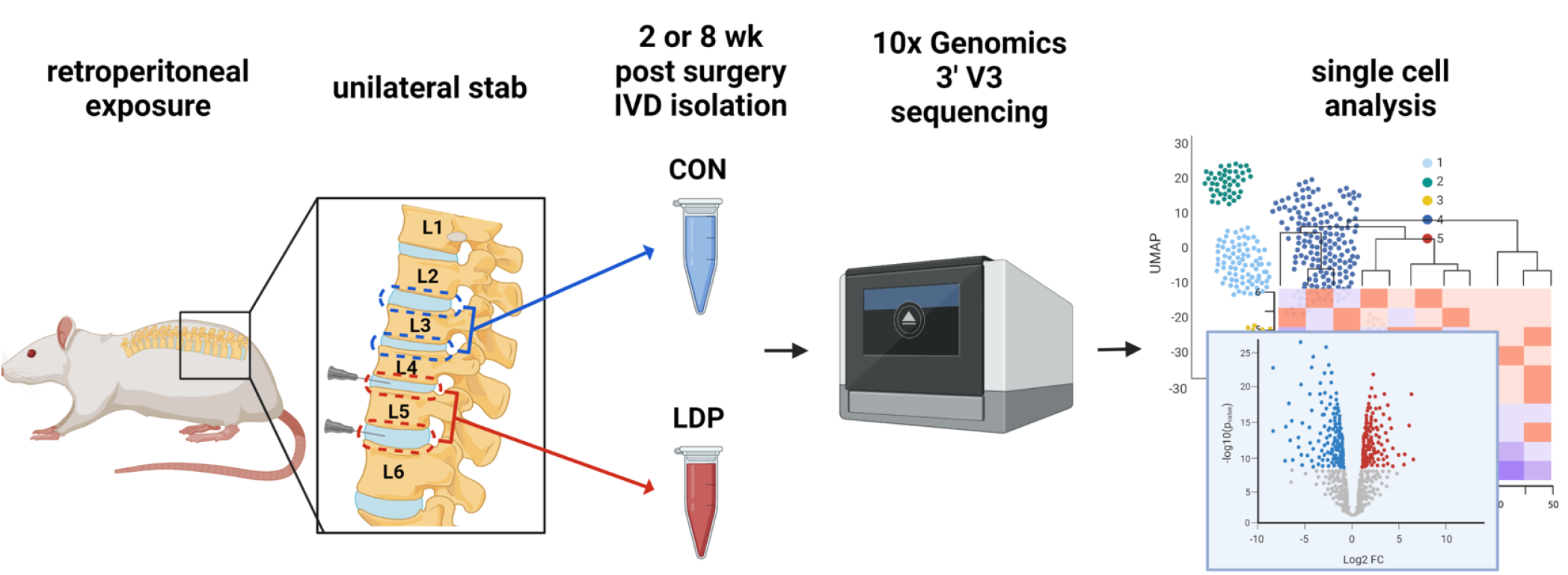
Flowchart depicting an overview of the single-cell RNA-seq experimental design.

### cDNA library generation and Single-cell RNA-sequencing (scRNA-seq) with data standardization

The input samples were submitted to the Washington University Genome Technology Access Center to obtain and sequence the cDNA libraries (10XGenomics, 3’v3.1; Illumina NovaSeq S4) according to established protocols. For CON and LDP samples, only cells with <10% mitochondrial features, gene counts of 500-3000 and UMI (unique molecular identifier) counts between 500-30,000 were determined to be of sufficient quality to include in the analyses (Partek Flow^TM^, Partek Inc). Under these selection criteria, the total number of cells from rat samples was 42560 and 51365 for CON and LDP at 2 weeks post-surgery, 27,064 and 28,293 for CON and LDP at 8 weeks post-surgery, respectively.

### Dimensionality reduction and unsupervised clustering

In order to identify resident cell populations, we first performed Principal Components Analysis (PCA) on pooled data from n=4 CON samples at 2 weeks, and then again for the 8 weeks samples. PCA was performed to reduce dimensionality of the data for CON at each timepoint with selection of optimum number of principal components after normalization; this process was repeated for the n=4 LDP samples at 2 and at 8 weeks. Cluster analysis was then performed with the iterative procedure of K-means clustering on CON cells at 2 weeks to identify optimal separation of cell clusters. A total of five major cell clusters were identified as providing for optimal separation of the data for CON samples. The percentage of cells mapped to each cluster was evaluated for each tissue sample at both 2 and 8 weeks post-surgery as a test of inter-subject variability and appropriateness of the clustering scheme. Annotated cluster data were visualized with uniform manifold approximation and projection (UMAP) analyses.

### Cell identification

The nomenclature classifying cell types within each identified cluster for CON samples at 2 weeks was based on a set of “marker genes” based on reports within the literature (see Table 1). For CON samples, we first validated the presence of known cell subsets based on prior scRNA-seq studies of intervertebral cells, endothelial cells, myeloid and lymphoid cells of the rat IVD [5, 15]. Violin plots of gene expression levels for all cells within each cluster were used to define the majority cell type in each cluster.

**Table 1.**
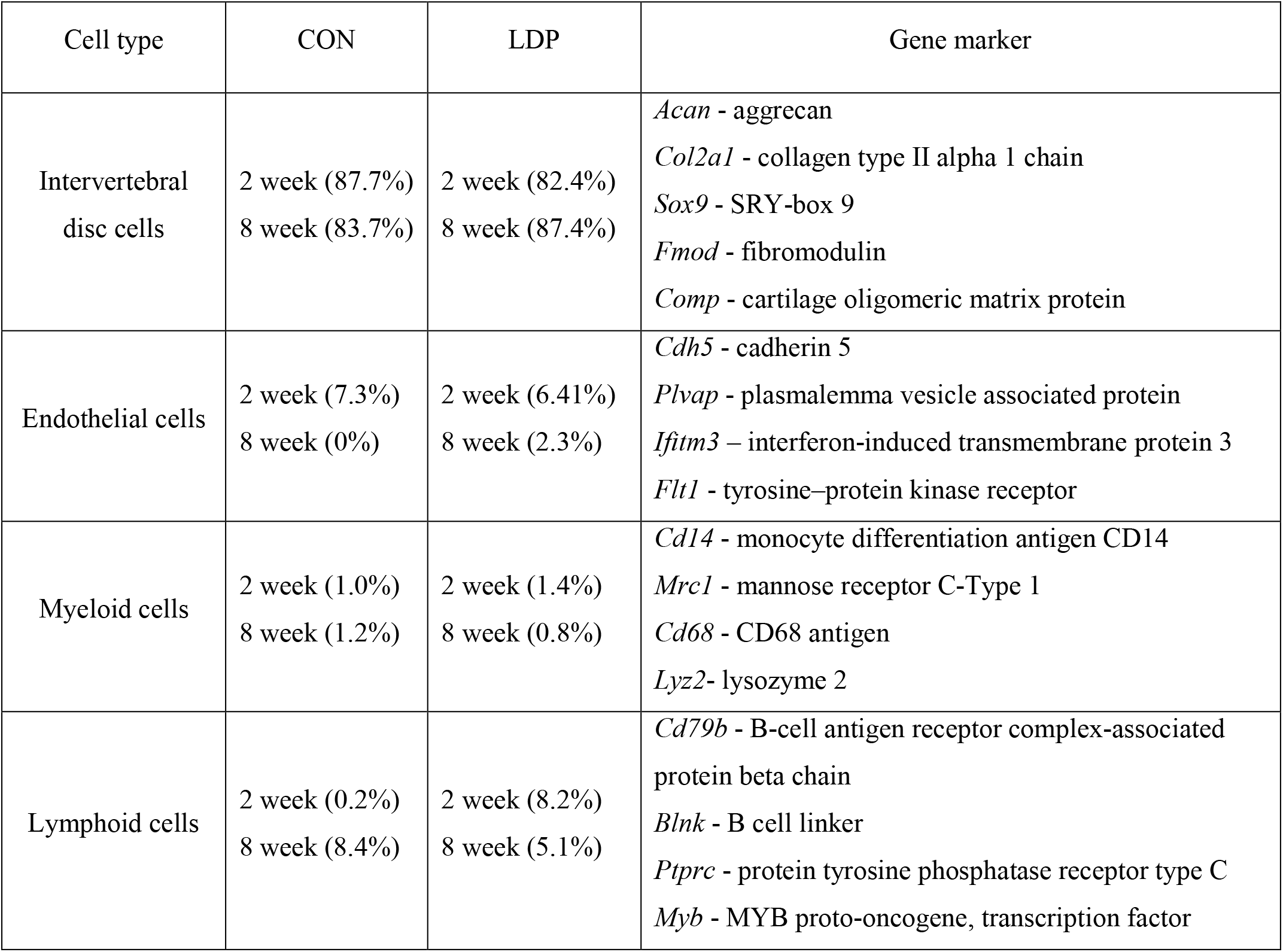
Genes examined for definition of cell sub-types in the IVD of rat samples (i.e., marker genes). Percentages are numbers of total cells assigned to each cluster averaged across four rats.^1^ CON – cells from control (unoperated) IVDs; LDP – cells from lumbar disc punctured IVDs at 2 or 8 weeks following surgery.

As a majority of cells in the CON populations were identified with a cluster named intervertebral disc cells, we sought to further sub-cluster this population in a post-hoc process separate from the data for cells mapped to the other clusters. IVD cells for CON samples at 2 weeks were subjected to PCA to select an optimal number of principal components, followed by K-means clustering on pooled data to identify further separation of cells in the intervertebral disc clusters based on the lowest Davies–Bouldin index.

The above-described process for clustering and sub-clustering was repeated for CON samples at 8 weeks post-surgery with no changes to cluster nomenclature nor marker genes. We sought to retain five major cell clusters for samples from the LDP population at both 2 and 8 weeks post-surgery, in order to directly compare cell numbers and gene expression levels against those of the CON population. In large part, the nomenclature for these five cell clusters did not differ from that of CON populations; however, the numbers of cells mapped to each cluster and gene expression for marker and novel genes varied substantially.

### Gene specific analysis

Differential gene expression analyses were done with the GSA toolbox (Partek Flow^TM^) to identify the differentially expressed genes within each rat sample between CON and LDP at each of 2 and 8 weeks post-surgery. The genes exhibiting the greatest fold-change were identified (log_2_ (fold change)) for each sample at 2 weeks or 8 weeks with a statistical significance of false discovery rate (FDR) p-value<0.001. This set of genes identified as most highly differentially expressed differed among samples with some overlap as described in results. Genes with the highest fold-difference between CON and LDP that are common across samples at 2 or 8 weeks are presented here.

### Gene set analyses

Gene set analyses were performed using the Gene Ontology resource (https://www.uniprot.org/help/gene_ontology) (UniProt Database) to predict the physiological roles and molecular processes of the upregulated gene networks. The top 50 differentially regulated genes for the respective cluster or subclusters were used to define the relevant cellular processes.

### Statistical analyses

Optimal separation of cells into a minimum number of distinct clusters was identified via the minimal Davies-Bouldin index, as indicated above for CON cells at 2 weeks and for IVD cells when separated into sub-clusters. Data for the magnitude of gene expression for cell-specific markers, e.g., genes defining B and T lymphocytes, were tested for differences between CON and LDP populations at each of 2 and 8 weeks post-surgery by the Student’s t-test or the Mann–Whitney test as appropriate (GraphPad Prism^TM^). Likewise, similar comparisons were made for the expression of *Ngf* and *Ngfr* (p75NTR) in matched sub-clusters between CON and LDP populations. Chi-squared tests were used to compare the proportions between matched cell clusters/subclusters across CON and LDP populations. Gene set enrichment analyses were performed by matching genes to defined pathways based on known cell functions and reporting overrepresented cell numbers in groups via a Fisher’s exact test or a (Partek Flow^TM^, Partek Inc).

## RESULTS

### Cell populations identified in IVD tissues from 2 weeks post-surgery

#### Samples in control IVD tissues 2 weeks post-surgery

Unsupervised K-means clustering was used for the CON samples to group the cells into clusters with equal variance. Results show that 5 clusters served to separate all cells into discrete populations. There were identified four named clusters: intervertebral disc cells, endothelial cells, myeloid, and lymphoid cells (Figure 2a); cluster 5 was annotated as “other cells” due to the lack of common identification markers. Identification of cell populations within each cluster was based on known gene markers that were pre-selected as described in methods (Figure 2b). The identified clusters had a distribution of cell counts that was very similar across samples for the four CON samples derived from four separate rats (Figure 2c). Each sample showed a similar cell distribution profile where intervertebral disc cells contributed to most of the cluster (∼87.7%), followed by endothelial cells (∼7.3%), myeloid (∼1.0%) and lymphoid cells (∼0.2%).

**Figure 2.**
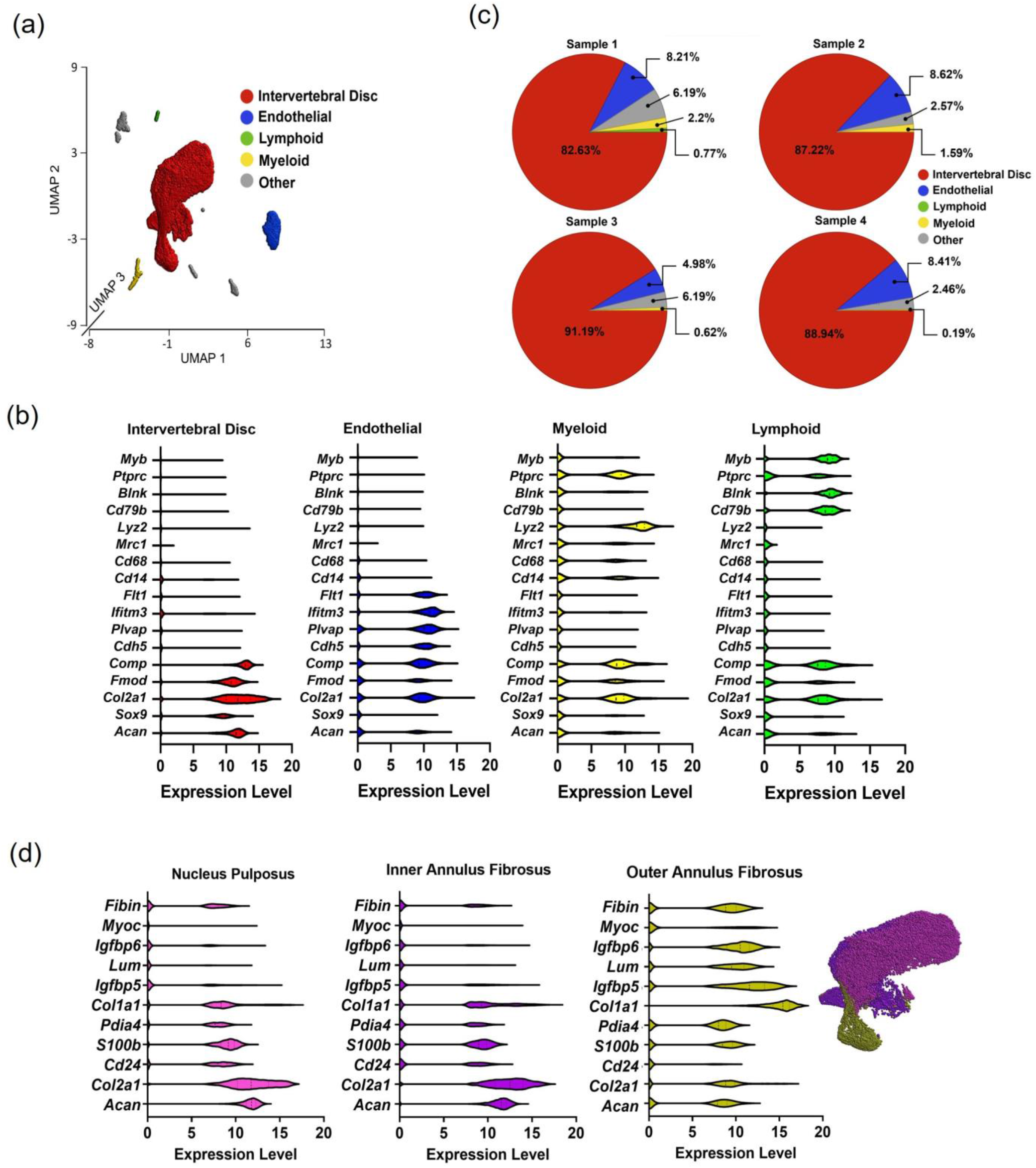
a) UMAP plot representation of five cell types within CON group at two weeks after surgery. b) Expression levels of marker genes within intervertebral disc cells, endothelial cells, myeloid, and lymphoid cells. c) Pie chart showing the distribution of each cell analyzed for the four rat samples in the CON group. d) The expression level of marker genes for three subclusters of intervertebral disc cells: nucleus pulposus, inner annulus fibrosus and outer annulus fibrosus.

The selected marker genes were chosen based on a priori knowledge for identifying the discrete cell clusters (Figure 2b). There was some commonality in extracellular matrix genes across clusters, primarily for *Col2a1* and *Comp*; this observation is consistent with the prevalent expression of these extracellular matrix proteins in different tissues and suggests that relying on non-extracellular matrix genes as differentiating markers may be preferable. Markers selected as lymphoid and myeloid markers proved to be effective in separating cells into clusters with more separation.

As more than 85% of all cells mapped to the “intervertebral disc cells” cluster, we sought to sub-cluster this population to identify the known cell populations. The IVD cell cluster was partitioned into three distinctively adjacent subclusters that were classified as nucleus pulposus (NP), inner annulus fibrosus (IAF), and outer annulus fibrosus (OAF) based on relative expression levels of classical IVD cell markers (Figure 2d). Subcluster 1 was identified to be NP cells based on a relatively higher expression for *Acan* and *Col2a1* while also expressing lower *Col1a1* compared to other IVD cells [8,29–32]. Additionally, these cells highly express *Cd24*, a classical marker for NP cells [13,33–35], previously identified markers of the chondrocyte lineage including S100B and a novel marker for bovine NP tissues, PDIA4 [19,36,37]. Cells mapping to the IAF cluster were similar in pattern of expression to that of NP cells, differing with modestly increased *Col1a1* expression and decreased presence of *Cd24*. Finally, OAF cells were identified based on the expression of a number of previously identified markers for this cell type including *Igfbp5*, *Igfbp6*, *Lum*, *Myoc*, and *Fibin* (Figure 2d) [8,19,29,31,32,38].

#### Samples in degenerated IVD tissues

Samples harvested from the LDP segments of the rats at 2 weeks post-surgery underwent unsupervised K-means clustering as applied to the CON group. Five clusters provided for optimal separation of the LDP cells based on known gene markers (Figure 3a, b). The IVD cluster contributed to greatest proportion of the cells (∼82.4%), followed by endothelial (∼6.4%), myeloid (1.4%) and lymphoid clusters (∼8.2%). The myeloid and lymphoid clusters exhibited the most notable difference 2 weeks after LDP (Figure 3c).

**Figure 3.**
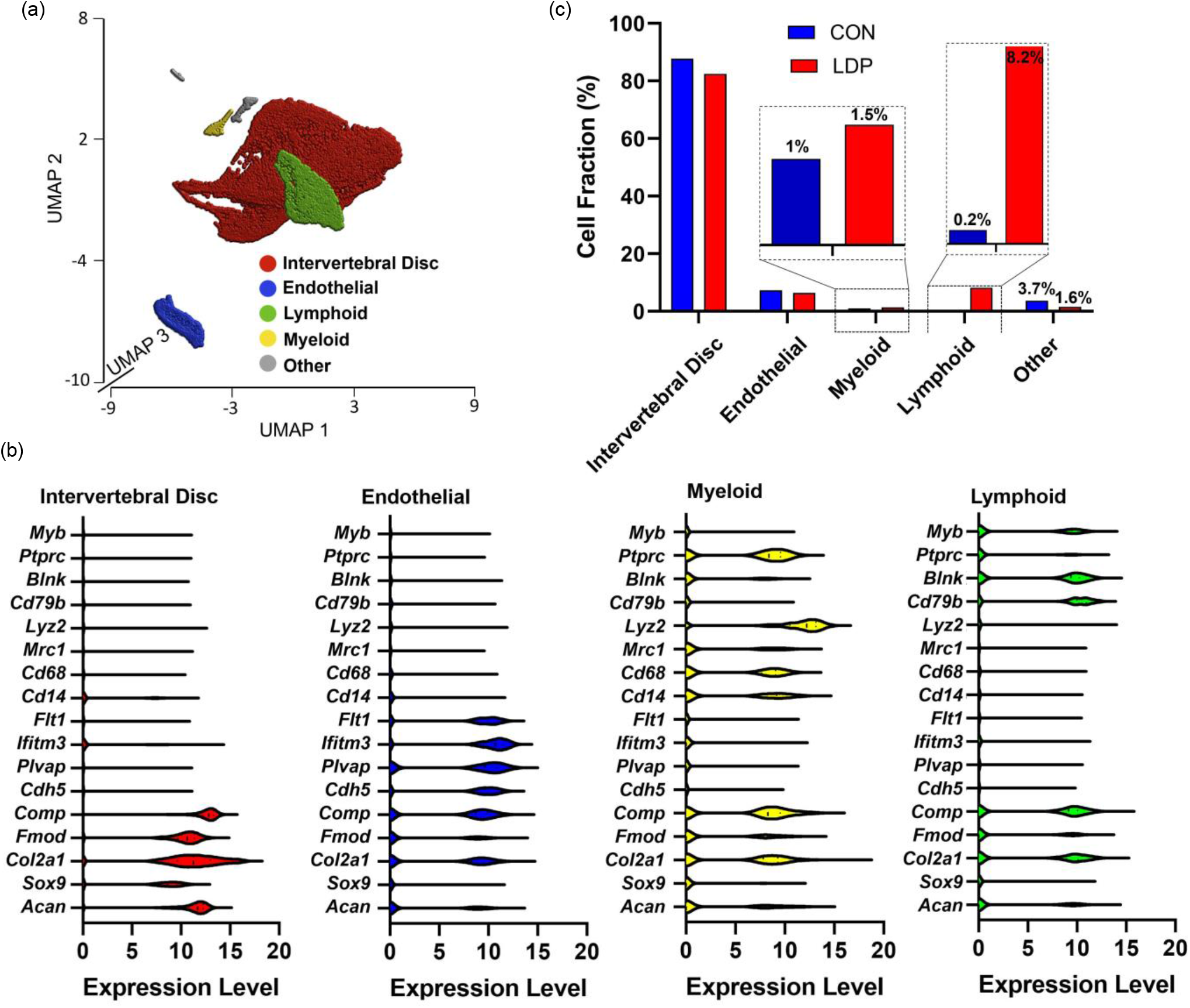
a) UMAP plot representation of five cell types within LDP group at two weeks after surgery. b) Expression level of marker genes within intervertebral disc cells, endothelial cells, myeloid, and lymphoid cells in LDP groups at two weeks after surgery. c) Percent of all analyzed cells mapped to each cell type for CON and LDP groups at two weeks after surgery. Inset provided to highlight differences in CON and LDP groups for immune cell populations)

#### Differential gene expression between LDP and CON samples on a subject-specific basis

To directly assess differences in mRNA levels for cells from the CON (L2-3 and L3-4 levels) and LDP (L4-5 and L5-6 levels) groups within each rat sample, we performed specimen-specific differential gene expression (Figure 4a, b). Differential gene analyses demonstrated that between 700-900 genes were more highly expressed in LDP cells at 2 weeks after surgery, as compared to their counterparts amongst the CON cell population (FDR p-value<0.001, Figure 4c). Analyses also showed between 250 and 1200 genes to be more highly expressed in CON cells as compared to the LDP cell population at 2 weeks after surgery (FDR p-value<0.001, Figure 4c). While there was variability in differentially expressed genes across samples, we identified a subset of 25 commonly upregulated genes in LDP populations of cells from two samples with the highest total numbers of sequenced cells (i.e. sample #3 and #4); expression levels for these genes against their cell identification are shown in the heat map in Figure 4d (representative results shown for sample #4). Of these 25 genes with upregulated expression levels in LDP compared to CON cells, the majority (22/25) were associated with cells of the lymphoid or myeloid cell clusters (Figure 4d).

**Figure 4.**
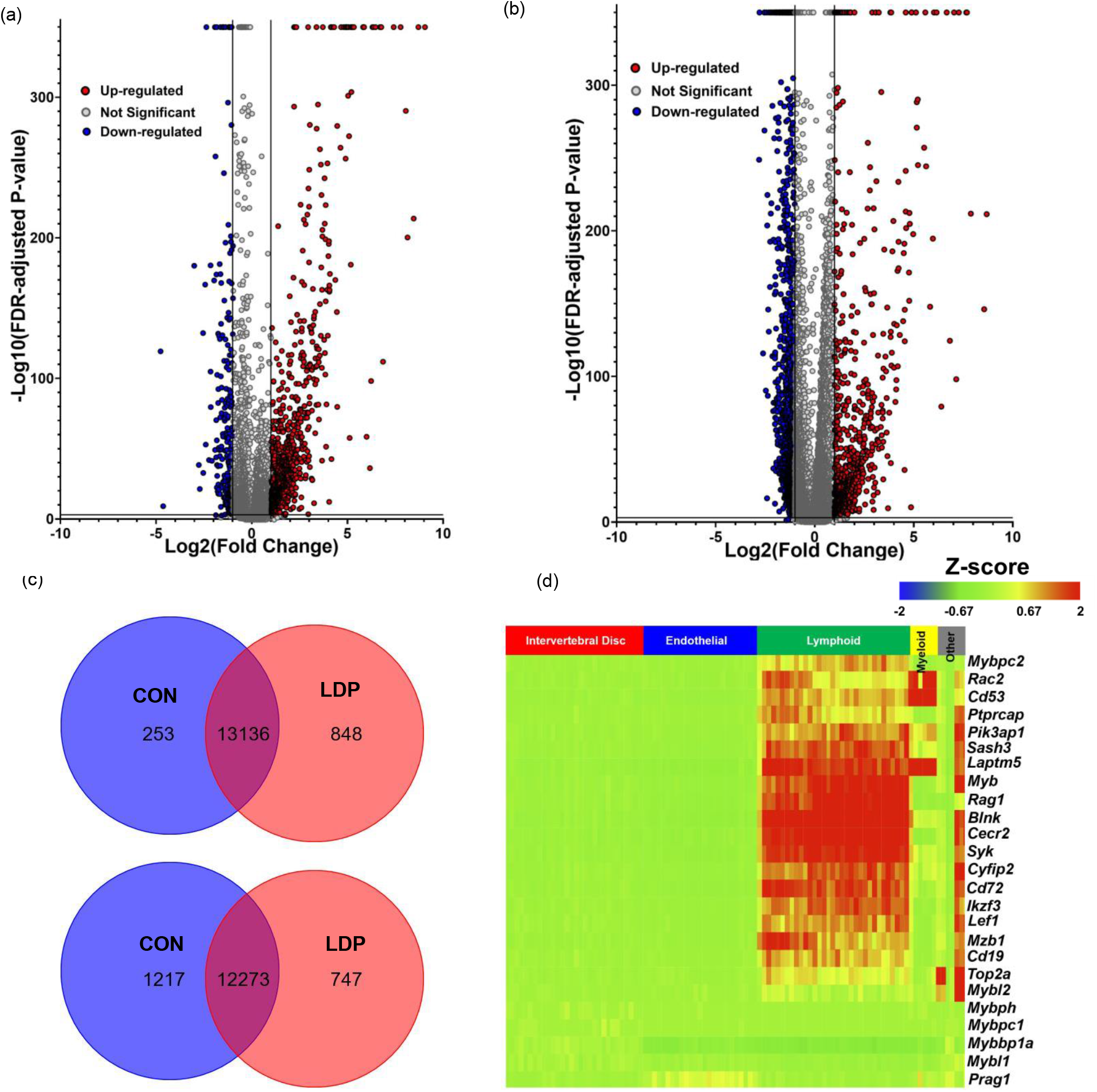
Volcano plot depicting specimen-specific differentially gene expression between CON and LDP cells at two weeks after surgery. Two representative samples are shown - a) sample 3 (left) and b) sample 4 (right) (Red dots represent genes expressed at higher levels in LDP group while blue dots represent genes with higher expression level in CON group) c) Venn diagram represents the total number of commonly upregulated and downregulated genes for LDP and CON cells in across multiple samples (i.e., samples 3 and 4 here). d) Heatmap showing the pattern for a subset of commonly upregulated genes with LDP compared to CON groups as a function of cell grouping (x-axis).

#### Immune cell involvement in differences between LDP and CON gene expression patterns

We determined the immune cell subtypes that were contributing to the upregulation genes due to LDP. First, there were significant increases in cell numbers mapping to the myeloid and lymphoid clusters for LDP samples as compared to CON samples at 2 weeks after surgery (p < 0.00001; Chi-square test), indicating an increase of infiltrating immune cells post injury (Figure 5a). Macrophages have been shown to infiltrate the herniated or injured disc in multiple species [23,39–44]. Accordingly, the number of cells expressing pan-macrophage markers (*Cd4*, *Cd14*, *Cd68*, and *Lyz2*), as well as their respective levels of expression, indicate increased macrophages and heightened activity (Figure 5b). *Cd4* is a pan-macrophage marker in rats [45], *Cd68* is involved in antigen presentation [46, 47], *Cd14* is essential for macrophage phagocytosis [48], and *Lyz2* is important for bacteriolysis [49]. In the LDP group, there were increased proportions of *Cd4*+ (p < 0.00001), *Cd14*+ (p < 0.05), *Cd68*+ (p < 0.00001) and *Lyz2*+ (p = 0.05) cells. Analysis of gene expression levels for each marker showed few differences in expression levels for these macrophage markers between CON and LDP groups, despite the higher numbers of cells expressing each marker in the LDP samples compared to CON (Figure 5b).

**Figure 5.**
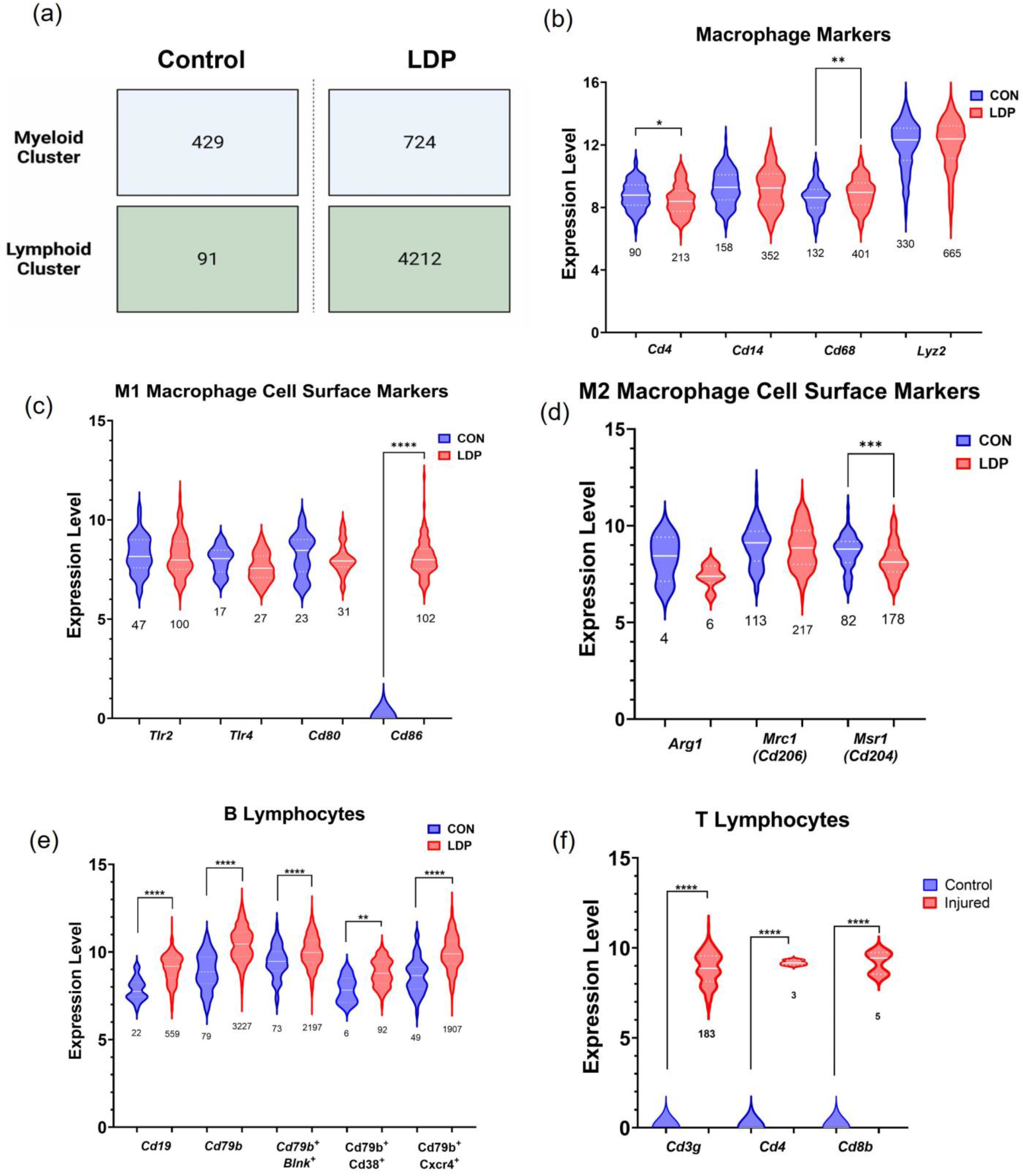
The identification of immune cell subtypes in the myeloid and lymphoid clusters. a) Diagram showing the differences in cell numbers mapped to the myeloid and lymphoid clusters between the control (CON) and degenerated (LDP) samples at two weeks after surgery. b) Pan-macrophage marker expression and cell count differences between CON and LDP groups were used to confirm the identification of cells in myeloid cluster. Macrophage polarization was assessed by identifying the presence of either c) M1 or d) M2 polarization cell surface markers, and the number of cells that expressed them. The presence of e) B and f) T lymphocytes was identified in the lymphoid cluster by measuring pan and subtype specific marker expression levels and cell count changes. Numbers underneath each plot denote the number of cells expressing each marker. A Student t-test or a Mann Whitney test was run to determine significance of marker expression levels. * = < 0.05, ** = <0.01, *** = <0.001, **** = <0.0001.

Macrophages can undergo polarization into the classically activated pro-inflammatory phenotype, M1, or the alternative anti-inflammatory phenotype, M2 [50, 51]. The number of cells expressing the canonical M1 cell surface markers *Tlr2*, *Tlr4* and *Cd80* were not different between groups, but the number of *Cd86*+ cells (p < 0.00001) and their expression were higher in the LDP group (Figure 5c) [51]. Despite the similarity in the number of cell expressing M2 polarization markers between LDP and CON groups, including *Arg1*, *Mrc1* (*Cd206*), *Mrs1* (*Cd204*), the expression levels were either indistinguishable (*Arg1*, *Cd206*) or lower (*Cd204*) compared with the CON group (Figure 5d) [51] [52].

The majority of the top 25 commonly upregulated genes were in the myeloid and lymphoid clusters, and gene analysis led to the intriguing discovery that over 80% percent of those genes were highly enriched in lymphoid cell lineages. Notably, B cell associated gene markers were the most represented group (40%: *Pika3p1*, *Rac2*, *Blnk*, *Syk*, *Ikzf3*, *Mzb1*, *Cd19*, *Mybl1*) while the T cell associated gene markers were slightly less prominent (25%: *Cyfip2*, Lef1, *Mybbp1a*, *Prag1*, *Myb*). Several gene markers were expressed in both T-and B-cells (35%: *Ptprcap*, *Sash3*, *Rag1*, *Top2a*, *Mybl2*, *Cd53*, *Laptm5*) (Figure 4d). B-lymphocyte specific markers included *Blnk*, *Rac2*, and *Pik3ap1*, which are involved in receptor signaling; *Syk*, *Cd72*, *Ikzf*, and *Cd19*, which are involved in differentiation and proliferation; and *Mzb1*, *Mybl1*, which are involved in antibody production and secretion.

Macrophages are capable of expressing cytokines that activate T-cells so that we examined T-cell specific markers for the incidence and genotype of T-cells. Notable T cell specific markers were *Cyfip2*, needed for T cell adhesion; *Lef1*, required for IL17A expressing T cell maturation; and *Myb*/*Mybbp1a*, which are involved in T cell differentiation. Five genes were highly expressed in both B and T lymphocytes and regulate a range of functions such as cell adhesion and migration, *Cd53*; antigen receptor signaling, *Ptprcap* and *Sash3*; and VDJ recombination, *Rag1* and *Top2a*.

The lymphoid cluster showed the largest difference in cell number between CON and LDP conditions, where the LDP samples had a cell count larger than 4.6x that of CON samples (p < 0.00001; Figure 5a). Therefore, we checked for the presence of B and T lymphocyte specific markers within the lymphoid clusters to confirm these immune cell types were present. B cell receptor and co receptor, *Cd79B* and *Cd19* in LDP samples, showed greater levels for gene expression compared to CON cells, and were expressed in the majority of the lymphoid cells (Figure 5e). *Blnk*, which plays a role in B cell maturation; *Cd38*, which is expressed on B cell progenitors and plasma cells [53, 54]; and *Cxcr4*, which plays a role in B cell activation and trafficking [55], all showed significantly greater gene expression levels in LDP samples concurrent with large increases in cell numbers between CON and LDP at 2 weeks after surgery (Figure 5e). *Blnk*+ (and *Cd38*+ cells were also higher in the LDP group. As for T lymphocytes, gene expression levels and numbers of cells expressing pan T cell markers, *Cd3g*, the T cell receptor, were both significantly increased (Figure 5f). Interestingly, only a small number of cells expressed *Cd4*, helper T cells, or *Cd8b*, cytotoxic T cells, emerged as identifiable in the LDP samples (Figure 5f). These data suggest that the majority of the cells in the lymphoid cluster are B and T lymphocytes with former being the largest subtype.

#### *Ngf* expression level variation in IVD cells in LDP populations

The number and marker expression of cells mapping to the IVD subclusters were evaluated between CON and LDP populations at 2 weeks after surgery. While few differences were observed in relative expression levels of *Ngf* across the subclusters for the LDP group, the number of cells expressing *Ngf* increased following injury (p < 0.00001; Figure 6a). In CON cells from the NP cell sub-cluster, just 7.9% of cells expressed *Ngf* which increased to 12.9% following injury (p < 0.00001). *Ngf* is believed to be important for promoting neurite infiltration into the otherwise avascular and aneural tissues of the NP [38]. In OAF cells, the incidence of *Ngf* expressing cells increased from 1.0% of cells in CON samples at 2 weeks after surgery to 3.8% of cells in LDP samples, a greater than three-fold increase (p < 0.00001). Additionally, the gene expression of NGF was significantly higher for all cells in the OAF subcluster of LDP samples (p<0.00001). Likewise, the expression of *Ngfr*, which encodes for p75NTR - a low affinity receptor for *Ngf*, was elevated in the OAF of LDP samples. Further, *Ngfr* was expressed in 2.3% of CON cells in the OAF sub-cluster increasing to 7.6% of cells expressing in the LDP injury group (p < 0.00001; Figure 6b).

**Figure 6.**
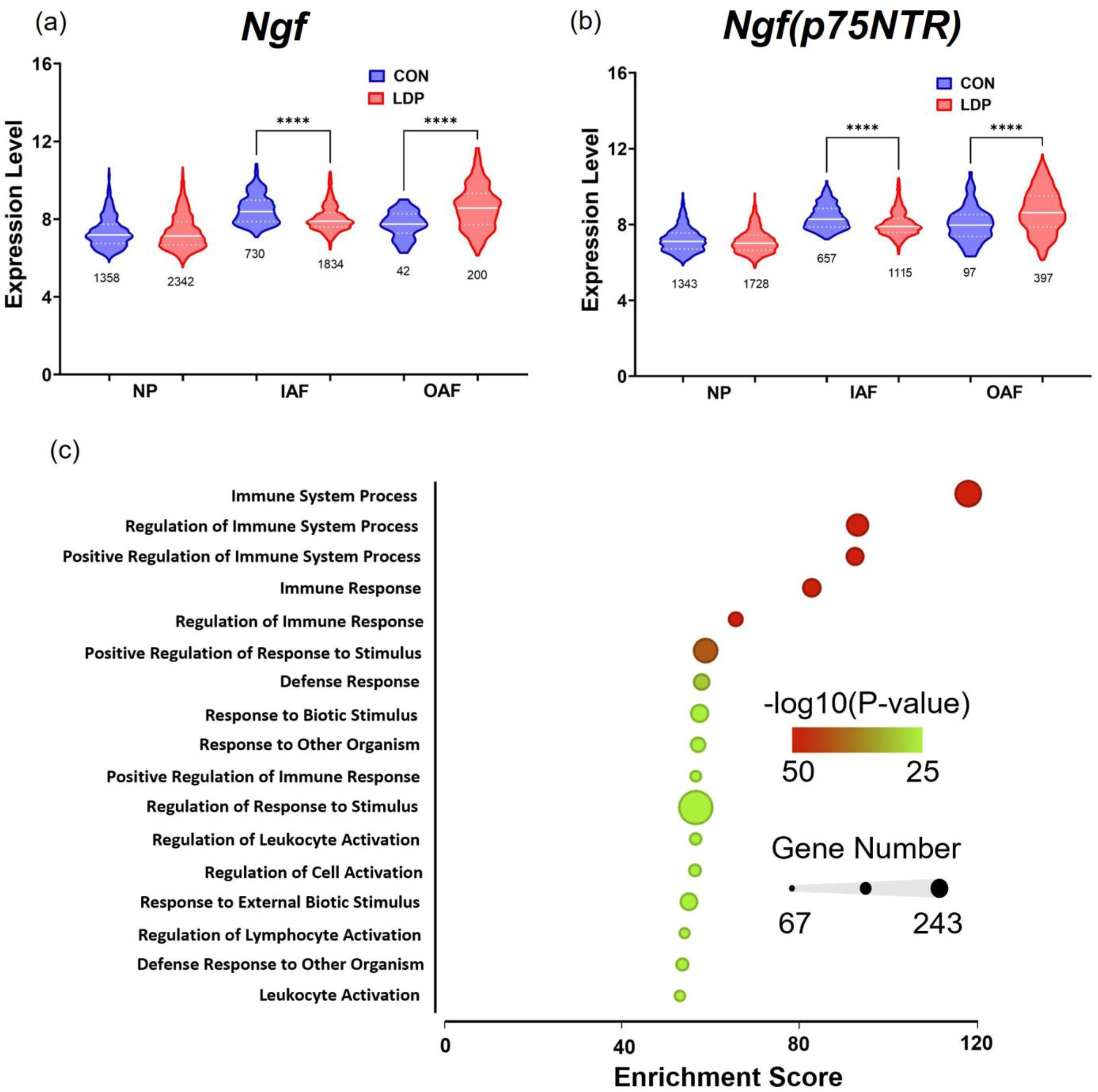
a) *Ngf* and b) *Ngfr* (p75NTR) expression and cell count counts for samples from CON and LDP populations at two weeks after surgery. Numbers underneath each plot denote the number of cells expressing each marker. c) Representative plot of gene set enrichment analysis of upregulated genes corresponding to one LDP sample. Gene ontology functions are ranked from top to bottom by enrichment score derived from Fisher’s exact test; The 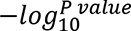 is represented by color where green denotes the highest level of statistical significance. The size of each node represents the number of genes in each gene ontology category.

The five most highly enriched processes identified by the Gene Set analyses were related to immune responses and associated regulatory processes (Figure 6c), consistent with the known infiltration of monocytes into the injured IVD post-surgical insult [40,56,57]. Genes associated with the top three regulated processes were upregulated in LDP samples and that are large drivers of the observed results are *Cd19*, *Cd37*, *Rag1*, *Blnk*, and *Myb*, as T- and B-cell associated marker genes.

### Cell populations identified in IVD tissues from 8 weeks post-surgery

#### Samples in control IVD tissues

As described for samples harvested at 2 weeks post-surgery, unsupervised K-means clustering was applied to the CON group at 8 weeks post-surgery. The 4 identified clusters had a distribution of cell counts that was very similar across samples for the four CON samples at 8 weeks post-surgery (Figure 7a); note that the absence of a 5th identified cluster was related to a cluster (previously “endothelial cell cluster”) with no mapped cells within. Violin plots of marker genes expression levels for all cells within each cluster were used to define the majority cell type in each cluster (Figure 1S). The IVD cells contributed to most of the cluster (∼83.7%) of all CON cells (8 weeks), followed by myeloid (∼1.2%) and lymphoid cells (∼8.4%). These differences from samples at 2 weeks post-surgery reflect a natural aging of the spine in the rat.

**Figure 7.**
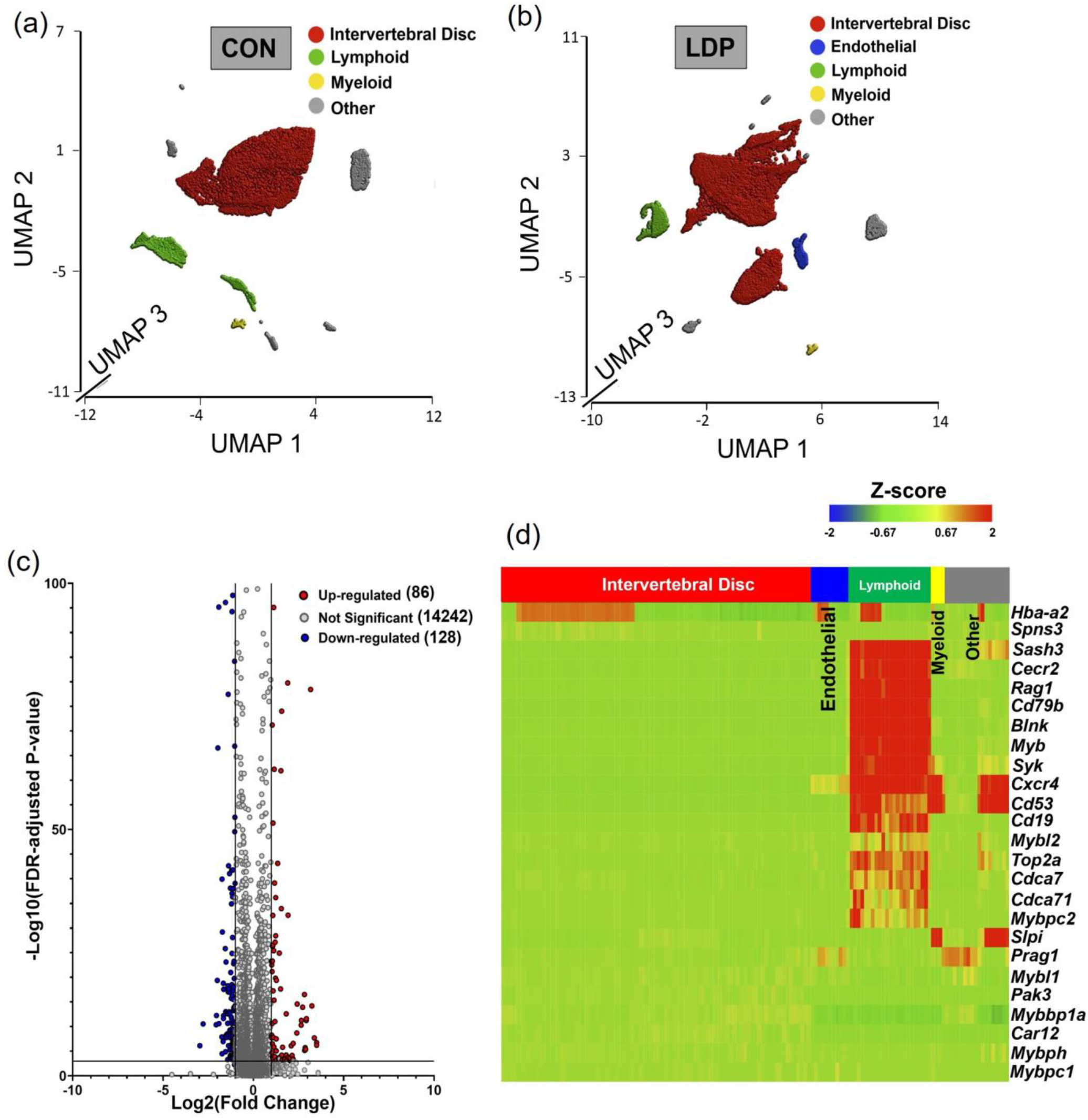
UMAP plot representation of all cell types within a) CON and b) LDP groups at 8 weeks after surgery. c) Volcano plot depicting genes differentially expressed between sample-specific LDP relative to CON populations for one sample at 8 weeks after surgery (Red dots represent genes expressed at higher levels in LDP while blue dots represent genes with higher expression level in CON group) d) Heatmap showing the pattern for a subset of commonly upregulated genes with LDP compared to CON groups at 8 weeks after surgery as a function of cell grouping (x-axis).

#### Samples in degenerated IVD tissues

Samples harvested from the LDP segments of the rats at 8 weeks post-surgery underwent unsupervised K-means clustering as described above, with the observation that five clusters provided for optimal separation of the data (Figure 7b). The IVD cells contributed to most of the cluster (∼87.4%), followed by endothelial cells (∼2.3%), myeloid (0.8%) and lymphoid cells (∼5.1%). Identification of cell populations within each cluster was based on known gene markers (Figure 2S). These differences in sample numbers reflect the remodeling response to LDP at 8 weeks after surgery.

#### Differential gene expression between LDP and CON samples on a subject-specific basis

Differential gene expression between CON and LDP cells at 8 weeks after surgery demonstrated that 86 genes were significantly upregulated in LDP compared to CON, while 128 genes were more highly expressed in CON than in LDP samples (representative plot shown for sample #8 in Figure 6c). A subset of 25 genes were identified as commonly upregulated in LDP compared to CON samples and used to generate a heat map, showing that the majority of genes identified were associated with cells of the immune system, including *Blnk* and *Cd79b* used to identify immune cells, along with *Myb*, *Rag1*, and *Cecr2*. These findings point to similarities in upregulated genes at the 2 and 8 weeks timepoints after LDP surgery; however, the identification of a lower number of upregulated genes and lesser fold effect changes suggest that the dramatic differences observed at 2 weeks after surgery in LDP cells may reflect an acute response to the injury.

The heatmap shows that the upregulated genes mostly belong to the lymphocyte cluster (Figure 7d). Therefore, we explore the presence of B and T lymphocyte specific markers within the lymphoid clusters. B cell receptor and co receptor, *Cd79b* and *Cd19* in LDP samples, showed greater levels of gene expression compared to CON cells, and were expressed in the majority of the lymphoid cells (Figure 8b). *Blnk*, *Cd38*, and *Cxcr4* all showed significantly greater gene expression levels in LDP samples concurrent with large increases in cell numbers between CON and LDP at 8 weeks after surgery (Figure 8b). As for T lymphocytes, the only gene expression is related to *Cd3g* with zero expression level for *Cd4* and *Cd8b* in both CON and LDP (Figure 8c). These data suggest that the lymphocytes cluster is almost composed of B cells by less sign of T cells with respect to 2 weeks.

**Figure 8.**
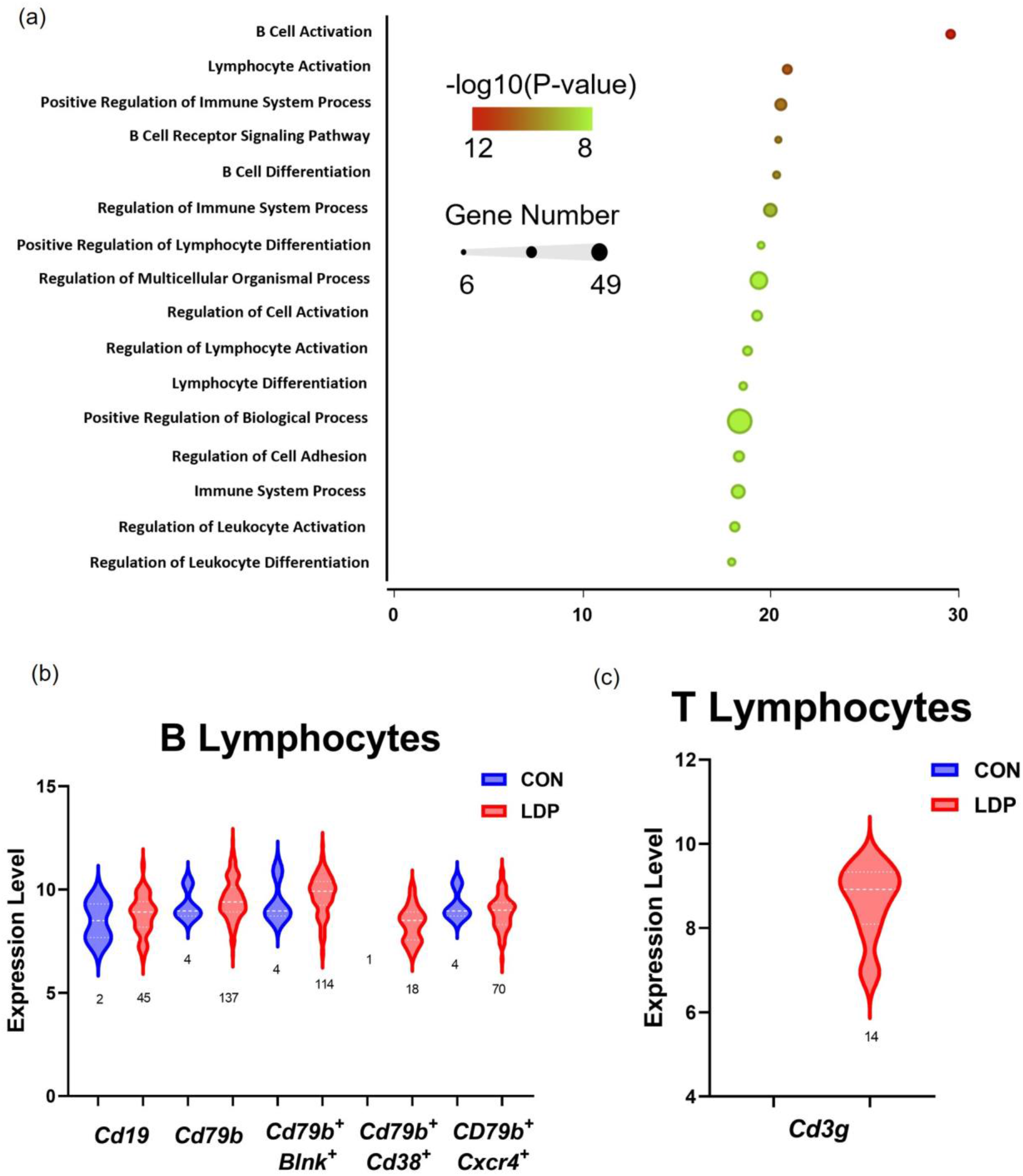
a) Representative plot of gene set enrichment analysis of upregulated genes corresponding to one LDP sample. Gene ontology functions are ranked from top to bottom by enrichment score derived from Fisher’s exact test; the 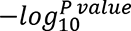 is represented by color where green denotes the highest level of statistical significance. The size of each node represents the number of genes in each gene ontology category. b) B and c) T lymphocytes were identified in the lymphoid cluster of one rat sample taken at 8 weeks after surgery (sample #8) by measuring pan and subtype specific marker expression levels and cell count changes. Numbers below each plot denote the number of cells expressing each marker.

Gene set analyses revealed the five most highly significant processes identified point to B cell activation, receptor signaling, and differentiation (Figure 8a). Genes associated with the top three regulated processes that were found to be upregulated in LDP samples and that are large drivers of the observed results are *Cd19*, *Cd7bB*, *Rag1*, *Blnk*, *Dppr4*, *Il7r*, *Lef1*, and *Myb*.

## DISCUSSION

Cells were isolated from the degenerated IVDs from a widely used rat model of lumbar disc puncture [24-28,58]. Single-cell RNA-seq was then performed on over 149,282 non-degenerate and degenerate IVD cells harvested from two timepoints following surgery to allow the progression of IVD degeneration. In brief, we identified four major cell types in the non-degenerate and degenerative IVD tissues of 2 weeks following surgery, based on the identification of genes previously identified as “classical” cell-specific markers [15]. For example, IVD cells closely clustered around classical “chondrocyte-like” markers including *Acan*, *Col2a1*, *Sox9*, *Comp* and *Fmod*. The cell classifications here in the master clustering and sub-clustering schemes for IVD cells from CON populations are consistent with a subset of those identified previously for NP cells; differences in cell identities were not noted between CON populations at 2 and 8 weeks following surgery, only differences in cell numbers mapping to each cluster.

Initial results that showed over 80% of all cells were identified as IVD cells at both 2 and 8 weeks, similar to a prior study showing that up to 99% of all sequenced cells mapped to a common transcriptomic profile [5]. The IVD cluster was then sub-clustered into NP, IAF, and OAF cells. For the markers chosen here for cell identification, analysis revealed the absence of any singular genes that are specific to each subcluster. Rather, IVD cell type identification requires a set of genes that are either more or less highly expressed when compared to their spatial counterparts. Classical markers of IVD cells such as *Acan*, *Col1a1*, and *CoL2a1* and some NP-specific markers such as *Cd24* were useful to support the sub-clustering scheme; however, expression levels exhibited a continuous distribution from inner to outer IVD. Prior studies have sometimes attempted to anatomically isolate NP tissues from adjacent inner AF prior to performing sc-RNA-seq analyses, yet yield the same outcome [5,20,22,59], indicating the spatial overlapping nature of these tissues and the transition between cell populations in the IVD exists in an continuum. Instead, we found additional gene markers including *Igfbp5*, *Igfbp6*, *Lum*, *Myoc*, and *Fibin* useful as supplements of the “classical” markers used to separate outer AF from NP populations within the IVD [5].

In the current study, effort was taken to evaluate sample-specific variability in the clustering results for the CON populations at each timepoint (e.g., Figure 2c). Studies that pool multiple tissue levels across animals to procure sufficient cells for a single sample for RNA-sequencing may be susceptible to one sample disproportionately driving the clustering outcome. While we had differences in cell yields between rats, the clustering scheme was valued for its ability to produce repeatable results across animals (e.g., 82-92% of all cells mapping to IVD cluster. Approaches that define correlation analyses for clustering schemes as used by Calio and co-workers [16] would be a useful addition to all sc-RNA-seq analyses that employ multiple tissues from multiple subjects.

With LDP, differences were observed in the numbers of cells associated with each identified cluster. We observed that the LDP samples contained 1.5 times more myeloid cells than CON tissue and that macrophages were the most prominent immune subtype present in the myeloid cluster (Figure 3c; Figure 5b). Macrophages have a remarkable range in cellular functions that macrophages can elicit during tissue injury and repair dependent upon macrophage polarization within the affected tissue [60]. M1-type macrophages are classically activated and differentiation is stimulated by IFNγ to secrete pro-inflammatory cytokines such as IL6, IL12 and TNFα. M2-type macrophages are alternatively activated macrophages, and their differentiation is stimulation by IL4 produced by Th2 T lymphocytes. Their roles are to secrete anti-inflammatory cytokines and factors such as Arginase 1, IL10, and aid in wound healing, tissue repair, and regeneration. The myeloid cluster were highly enriched for both M1 macrophages and M2 macrophages at 2 weeks post IVD injury (Figure 5 c,d). While both M1 and M2 macrophage markers are present, there is significantly higher expression for *Cd86* (M1) and significantly lower expression for *Arg1* (M2) genes, suggesting a shift towards M1, or pro-inflammatory state with LDP at 2 weeks post injury. This shift away from the M2 phenotype is consistent with the observations in the human degenerate nucleus pulposus cells that exhibited declining M2 proportion with increasing degeneration severity. Single cell transcriptomic analyses of the human IVD cells show also the increasing activation of immune recruitment pathways during degeneration, demonstrating the robust interactions between the immune system and the degenerating IVD.

Our study here also identified the increased activation of B-and T-lymphocytes two-weeks into the degenerative process. Though T-lymphocytes have been found in degenerating human IVDs [5], the presence of B-lymphocytes, which accounts for the majority of the highly expressed genes in the LDP population (e.g., *Cd72*, *Blnk*, *Syk* and *Rag1*) has yet to be described during degeneration (Figure 5e, f). The involvement of immune cells and hematopoietic cells in the LDP tissue suggest that cell infiltration and differentiation were promoted by the surgical stab injury [14]. The role of lymphocytes after tissue injury had not been studied extensively; recent studies in bone fracture healing show clear roles for lymphocytes in mediating the tissue response after injury [61]. In bone fracture healing, lymphocytes were observed to infiltrate the injury site in a temporal fashion where they initially infiltrate the tissue early during the inflammation process, by 3 days post injury, then leave and return after the peak inflammation window has passed and tissue proliferation has begun, around 14 days post injury [61]. Lymphocytes function in the inflammation process by recruiting antigen presenting cells, secreting cytokines to help mediate the inflammatory response, and secreting growth factors to aid in regeneration [61, 62]. We also observed that lymphocytes persisted up to 8 weeks post injury (Figure 8a,b; Table 1). B-lymphocytes act as antigen presenting cells and dampening the inflammatory response by promoting Treg survival and secreting IL10 [62]. The lymphoid cluster contained *Cd79b*/*Cd38* positive mature B-lymphocytes [54], and the presence of *Cd79b*/*Cxcr4* positive B-lymphocytes indicate activation and trafficking (Figure 5e,f) [55]. T-lymphocytes, in contrast, modulate antigen presenting macrophages, and regulate the inflammatory response with specific subtypes like *Cd4* positive Tregs [63]. *Cd4* positive T-lymphocytes are associated with tissue repair after injury due to eliciting immunomodulatory effects, while cytotoxic *Cd8* positive T cells have been shown to have a negative effect on repair [61]. We observe *Cd3g* broadly expressed in T-lymphocytes, *Cd3g/Cd4* expression in T-lymphocytes helper cells, and *Cd3g/Cd8* expression in cytotoxic T-lymphocytes (Figure 5e,f). Our data here reveal a novel role for B-lymphocytes in concert with T-lymphocytes as part of the immune cascade during degeneration that persists in both our model here and in humans. Moreover, the immune markers, such as *Myb*, *Blnk* and *Cd53*, remain similar over time for CON and LDP, suggesting that this is a sustained, progressive response due to degeneration rather than the acute trauma of the injury.

In addition to the described differences in cell number and gene expression between cells within the clusters of control and degenerative IVDs, we observed changes in *Ngf* levels and expression pattern for IVD cell sub-clusters with IVD degeneration. Both *Ngf* and *Ngfr* (encoding for p75NTR) were increased in expression levels in OAF cells, and the number of OAF cells expressing *Ngfr* was also increased. Additionally, NGF is pro-angiogenic that upregulates VEGF and indirectly recruits endothelial cells [64]. Sustained NGF sensitizes innervating primary afferent neurons to produce hyperalgesia [65] via upregulated NGF receptor expression in ganglion neurons [66], and promote expression of substance P, calcitonin gene-related peptide, and neuronal ion channels that may lead to long-term adaptive effects on nociceptors [67, 68]. Moreover, NGF binding to its receptor can regulate collateral sprouting of sympathetic fibers to dorsal root ganglion neurons that may relate to maintenance of sensitization or chronic pain. Indeed, a prior study of this LDP model in the rat revealed elevated protein expression for the NGF receptors TrkA and p75NTR in the innervating dorsal root ganglion neurons [58]. Secreted NGF can also drive growth and differentiation of both B- and T-lymphocytes [69, 70] in a pattern consistent with the elevated expression of these cell populations in the LDP samples.

A limitation of the current study is the absence of PCR confirmation of identified targets as performed in some prior studies [5,15,22]. As our interest was on numerically minor cell types involved in IVD degeneration, rather than phenotyping cells derived from discrete regions of the IVD, both immunolabeling and bulk-PCR proved difficult with remaining sample-specific tissues. Future work would be needed to design a surgical approach that made use of more lumbar levels, or other strategies such as flow cytometry, to support collection of tissues and cells so that bulk-PCR or immunolabeling could be used to confirm the current findings. As is, the clear corroboration of clustering schemes that supported use of “classical” cell-specific markers in the rat as used by Wang and co-workers [15], and the consistency observed in clustering results across four distinct rat subjects provides some measure of confidence and consistency to support the results provided here.

In summary, this study of single-cell RNA-seq in a rat model of IVD degeneration demonstrated changes in the numbers of cells associated with immune cell populations, with increased infiltration of B-lymphocyte and macrophage subsets into the IVD, particularly in early degeneration. While the surgical stab of the IVD undoubtedly provoked the release of soluble factors [71] that support endothelial cell infiltration, neurite invasion and the activation of infiltrating monocytes that lead to neuronal sensitization and the identification of B- and T-lymphocytes as shown here, the results also point to a region-specific increase in the number of cells expressing *Ngf* and *Ngf*r (p75NTR) and associated transcriptional expression that may be one of the key findings that support the development of progressive IVD degeneration following stab injury to the IVD in this model and to annulus injury in the human.

## ACKNOWLEDGEMENTS

Funding was received from the National Institutes of Health (R01AR070975, R01AR078776, R01074441, T32 EB028092). The Orthopedic Research and Education Foundation (resident research grant P20-03413), the National Science Foundation (Graduate Research Fellowship No. DGE-1745038), and the Rita Levi-Montalcini Postdoctoral Fellowship in Regenerative Medicine. We also acknowledge the Genome Technology Access Center (GTAC) at Washington University School of Medicine for help with genomic analysis (NIH P30 CA91842, NIH UL1TR002345). Figure 1 was created with Biorender.com.

## Author contribution

M.R, S.C, G.E, D.P and F.L contributed to study design, performed research, and analyzed data. D.P and L.J preformed experimental test. L.A.S, J.J.S and S.Y.T designed the research study and contributed to data interpretation. M.R, S.C, D.P, J.J.S, S.Y.T and L.A.S wrote the manuscript and contributed to data interpretation. All co-authors reviewed and revised the manuscript.

